# Real-time Detection of Driver’s Movement Intention in Response to Traffic Lights

**DOI:** 10.1101/443390

**Authors:** Zahra Khaliliardali, Ricardo Chavarriaga, Huaijian Zhang, Lucian A. Gheorghe, Serafeim Perdikis, José del R. Millán

## Abstract

Movements are preceded by certain brain states that can be captured through various neuroimaging techniques. Brain-Computer Interfaces can be designed to detect the movement intention brain state during driving, which could be beneficial in improving the interaction between a smart car and its driver, by providing assistance in-line with the driver’s intention. In this paper, we present an Electroencephalogram based decoder of such brain states preceding movements performed in response to traffic lights in two experiments: in a car simulator and a real car. The results of both experiments (N=10: car simulator, N=8: real car) confirm the presence of anticipatory Slow Cortical Potentials in response to traffic lights for accelerating and braking actions. Single-trial classification performance exhibits an Area Under the Curve (AUC) of 0.71±0.14 for accelerating and 0.75±0.13 for braking. The AUC for the real car experiment are 0.63±0.07 and 0.64±0.13 for accelerating and braking respectively. Moreover, we evaluated the performance of real-time decoding of the intention to brake during online experiments only in the car simulator, yielding an average accuracy of 0.64±0.1. This paper confirm the existence of the anticipatory slow cortical potentials and the feasibility of single-trial detection these potentials in driving scenarios.

## 1 Introduction

Non-medical applications for Brain-Computer Interfaces (BCI) are increasingly being explored, in particular for real-time monitoring of mental states, such as attention, performance capability, emotion, etc.^3,4^. Along this line, the possibility of decoding brain activity, measured with electroencephalography (EEG), to improve driving assistance systems has recently attracted considerable interest^11,13,14,29^. In this approach, the objective is not to ‘control a car with a BCI’ but rather to predict the driver’s intentions^11,13,14,29^. ‘in order to help the driving assistance system of a smart car’.

Nowadays vehicles are increasingly being equipped with technologies that provide environmental information, which may warn about critical situations during driving and eventually help to reduce the driver’s physical and cognitive workload^12,26^. By including the BCI, the assistance will not only be in-line with the situations on the road (using information from in-car sensors) but also, and more importantly, aligned with the driver’s intentions (mediated by the BCI).

During driving, we are constantly engaged in responding to external events. At a traffic light, the driver *anticipates* the appearance of the next color and actively prepares to perform the right action (i.e., accelerate or brake) at the proper moment. In case of an inattentive driver, unaware of the need to brake, the BCI could detect the absence of neural correlates of the anticipatory process. The driving assistance could thus provide feedback to the driver and eventually initiate the braking action gently, so that the driver becomes aware of the situation and has the chance of braking by themselves. This kind of assistance would prevent an automatic emergency braking at the last moment, which may result in a negative surprise and an unpleasant experience for the driver who could feel under the control of the smart car, rather than being in control of it. If the driver is aware of the color change of the traffic light and has the intention to brake, the BCI detects the presence of an anticipatory brain potential and an unnecessary automatic braking is avoided. Still, further driving assistance could be provided by facilitating the driver’s intended action.

At a traffic light the yellow light is a warning stimulus that predicts the appearance of the upcoming ‘Red’ stimulus, upon which the driver needs to perform an action (braking). Classical psychophysical paradigms to study anticipation have shown a slow negative deflection within the interval between the warning and the imperative stimuli called Contingent Negative Variation (CNV)^27^, which is a type of Slow Cortical Potential (SCP). The SCPs are defined as slow negative deflections observed in EEG lasting from 300 ms to several seconds with magnitudes up to 40*μ* V during specific cognitive tasks^2^.

The main goal of this paper is to investigate the neural signatures of the driver’s anticipatory behavior in response to traffic lights. In previous work^13^, we have investigated the neural correlates of anticipatory behavior with a count-down paradigm in a driving simulator. The results demonstrated the possibility of predicting the driver’s movement intention through anticipatory behavior in offline analysis. Here, we extended our approach to include (i) a more realistic driving scenario (traffic lights) in simulated and real car driving, (ii) assessment of the real-time detection of anticipatory brain potentials during online experiments in the car simulator, and (iii) preliminary assessment of the effect of providing feedback on the anticipatory behavior.

## 2 Materials and Methods

We have conducted two experiments in order to investigate the neural signature of the driver’s anticipatory brain state while driving in the car simulator and in the real car (both provided by Nissan Motor Co). The protocol for both experiments was approved by the institutional ethical committee and all subjects provided written informed consent.

For both experiments, EEG was acquired at a sampling rate of 2048 Hz using 64 electrodes according to the extended 10/20 international system (Figure 1) along with two EMG electrodes (Biosemi ActiveTwo, The Netherlands). Electrodes were placed using an elastic cap fitted with electrode holders. Conductive electrolytic gel had to be applied prior to the beginning of the experiment but no re-application was needed afterwards. The EMG signal was recorded using one set of bipolar electrodes placed on the *tibialis anterior* muscle of the subjects’ right leg^13^.

**Figure 1:**
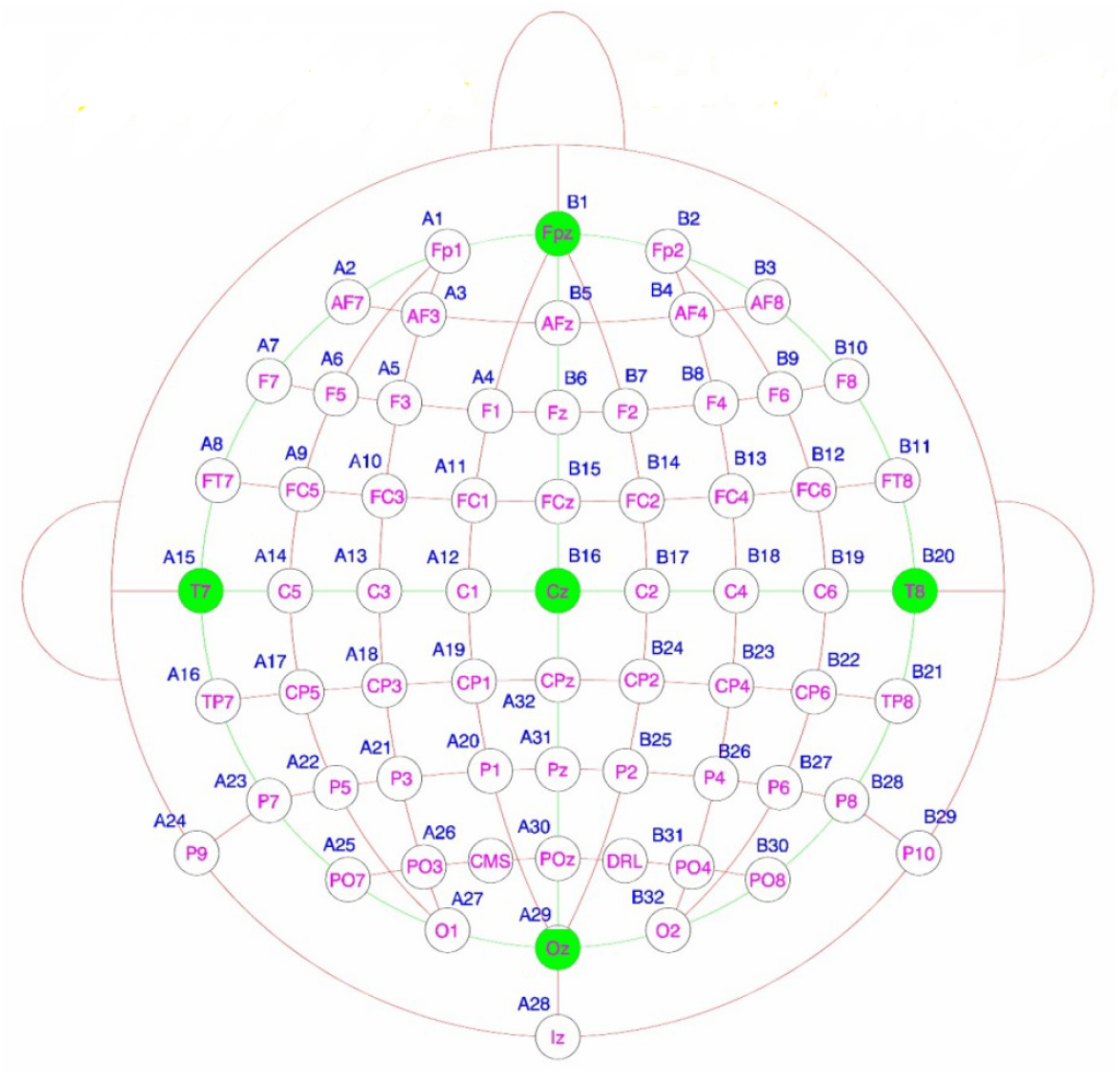
64 EEG Electrodes layout (extended 10-20 international).

For Experiment-1, which was held in the car simulator, comprised both offline and online experimental sessions, as follows. **Offline** sessions refer to the recordings where the analysis has been performed after the experiment (the signal processing and classification pipeline was implemented over recorded data). In the **online** sessions, the signal processing and classification pipeline was implemented in real-time. The online experiments entail the ‘real-time’ decoding of anticipatory brain potentials and allows to provide some feedback of the real-time detection to the user (i.e., closed loop).

The chain processing steps for the offline and online session are exactly the same for the feature extraction and classification steps. However, those processing chains differ slightly in the pre-processing steps, as explained in detail in Section 2.1.2 (Offline analysis and online analysis, respectively). For experiment-2, which was held in the real car, we have only conducted offline sessions. The processing chain of this experiment is explained in detail in Section 2.2.2. Both experiments comprised several recording sessions over different days. We firstly demonstrate the results of the post-hoc analysis of the data recorded in Day-1 of each experiment (offline sessions): i. Grand averages to confirm the presence of the anticipatory SCPs during the color changes of the traffic lights. ii. The single-trial analysis represented in ROCs. The results of online experiment-1 is shown in Table 3, which reports the accuracy of real-time classification in the subsequent recording days.

For both experiments, we performed a post-hoc analysis to confirm the presence of the anticipatory SCPs during the color changes of the traffic lights and report grand average Event-Related Potentials (ERPs) on the offline sessions. We also assessed the singletrial analysis decoding performance and report it using the Area Under the Curve (AUC) in the Receiver Operating Characteristics (ROC) space. For Experiment-1, the performance of the online sessions is reported in terms of the accuracy of real-time classification.

### 2.1 Experiment-1: Car simulator

#### 2.1.1 Setup and protocol

Ten healthy subjects (1 female, average age 23.5±1.4 yrs) participated in the experiments with a custom-made driving simulator (see Figure 2.a) over 3 days. All subjects possessed a valid driving license and had normal or corrected-to-normal vision. None of the subjects were color blind. They sat in the driver’s chair of the simulator in front of three 3D monitors (27 inches, 50 cm distance to the subject’s eyes, no need for wearing 3D glasses). We used the open source *VDrift* driving game^15^ and we designed a 3D virtual environment using the *Blender* software^1^ simulating a city with two main streets and five smaller streets perpendicular to them, yielding 10 intersections. Traffic lights were placed besides the road at each junction (see Figure 2.b).

**Figure 2:**
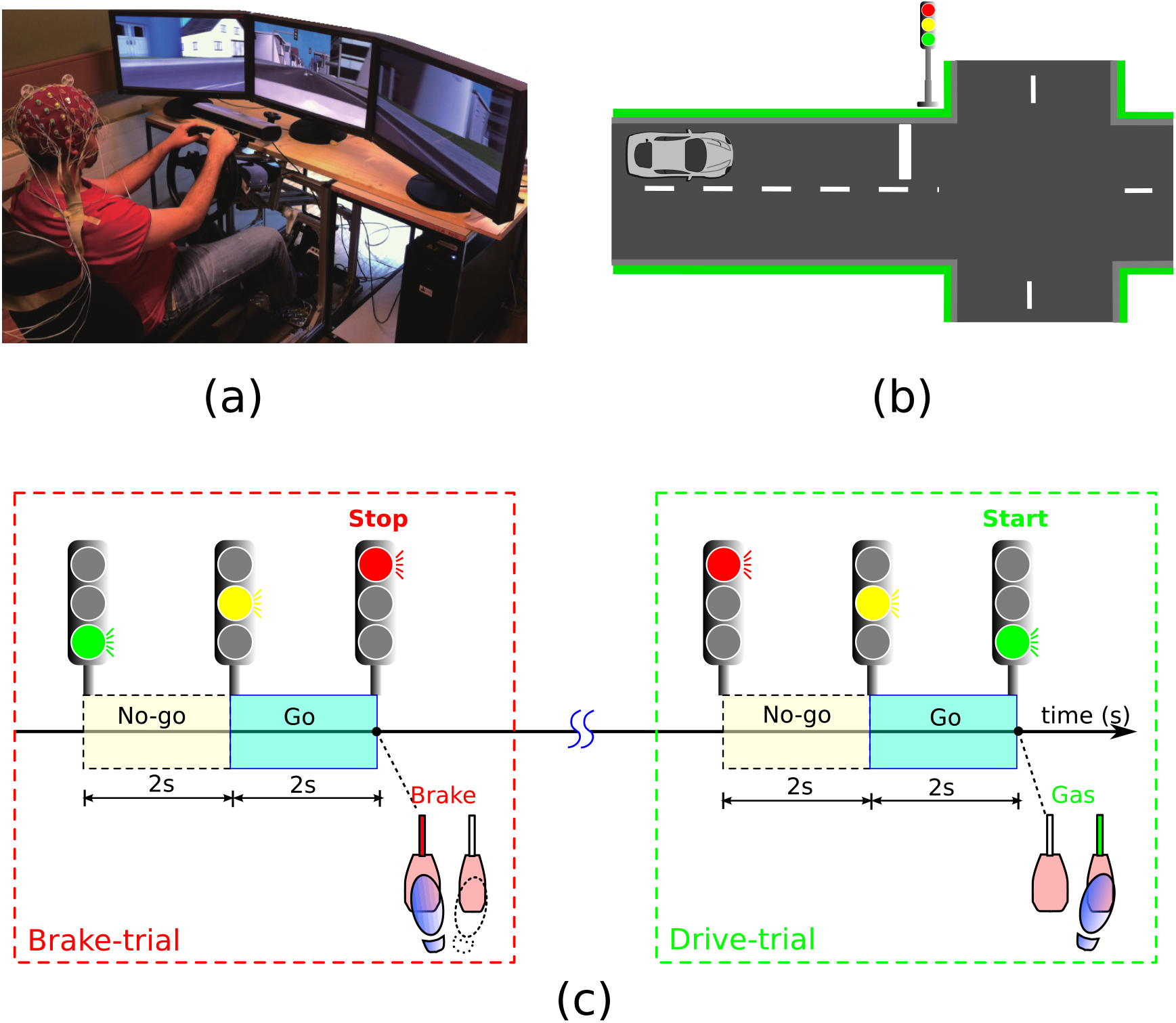
Experiment-1: (a) Experimental setup in the car simulator with three 3D monitors, (b) Illustration of a sample junction in which the car is approaching the junction with a traffic light in ‘Red’. (c) The time-line of the protocol: the traffic lights turning from ‘Green’, to ‘Yellow’, to ‘Red’ followed by braking after the ‘Red’ light, named *Brake* trials; this is followed by a waiting period for approximately 12 s; then, the next sequence of traffic lights with ‘Red’, to ‘Yellow’, to ‘Green’ corresponds to the *Drive* trials. Each type of trial contains one *Go* and one *No-go* epoch.

Subjects had to drive the virtual automatic car through the city, using the steering wheel, accelerator and brake pedals. To reduce signal contamination, subjects were instructed to fixate their gaze on a cross point at the center of the screen and minimize facial or head movements during the appearance of the stimuli (traffic light changes). They were asked to maintain a fixed speed of 80 Km/h. The simulated environment was programmed so that when the car approached a junction, the traffic light changed from ‘Green’, to ‘Yellow’, to ‘Red’. Subjects were instructed to press the brake pedal only at the onset of the ‘Red’. After the virtual car stopped, the driver had to wait for approximately 12 s for the next round of color changes of the traffic light (‘Red’ to ‘Yellow’ to ‘Green’). The duration of each color was set to 2 s. To make the subjects more involved, in 20% of the cases (randomly distributed) the traffic light remained ‘Green’ when the car approached the intersection (the driver was not required to stop).

This design allowed us to investigate the anticipatory brain potentials during driving, and more specifically, to test the difference between predictable future events in which the subject is not required to immediately perform an action (onset of ‘Yellow’ light), and imperative ones (light changes to ‘Red’ or ‘Green’), in which the subject is supposed to perform an immediate action. We defined two types of trials (see Figure 2.c): *Drive* (comprised the time interval from the ‘Red’ to ‘Green’ light) and *Brake* trials (the time interval between the onset of the ‘Green’ to ‘Yellow’ to ‘Red’ light). Each type of trial contained one *No-go* epoch and one *Go* epoch. The former corresponds to the interval between the change of ‘Green’ or ‘Red’ to ‘Yellow’, in which subjects were not supposed to perform any action after the cue, while the latter corresponds to the interval between ‘Yellow’ and ‘Red’ or ‘Green’, in which subjects were supposed to perform a specific action.

The experiment comprised between 2 and 3 recording sessions, as explained below. Each day of recording consisted of 4 sessions, and each session consisted of 4 runs. Each run lasted around 15-20 minutes and contained an average of 77±8 and 78±10 *Drive* and *Brake* trials, respectively, across all recording 3 days. In all sessions, no feedback (NF) was provided for the *Drive* trials. In contrast the subject Reaction Time (RT) for *Brake* trials was provided on the screen to the subject as a behavioral feedback 2s after the onset of the ‘Red’ light. RT was calculated as the difference between the timing of the onset of the ‘Red’ light and the pressing the brake pedal (see Table 1 for the details of feedback over the different sessions).

**Table 1.** Details of the type of feedback in the different recording sessions for *Drive* and *Brake* trials. NF: No Feedback, RT: Reaction Time; behavioral feedback; BCI: the output of the real-time EEG classification.

|  | Day-1 | Day-2 | Day-3 |
| --- | --- | --- | --- |
| *Drive* | NF | NF | NF |
| *Brake* | RT | RT+BCI | RT+BCI |

Only for *Brake* trials, using the data of Day-1, a classifier was trained and used for online experiments. Those subjects for whom the classifier yield classification performances above random performed online sessions in the next recording day (Day-2). Five of them (S1, S2, S3, S4, and S5) also performed an additional online (Day-3). The training classification performance on Day-1 was not satisfactory for 3 of the subjects (S7, S9, and S10). For S7 and S9, the training classification performance was very close to the chance level. This was presumably due to artifact contamination. Therefore, we repeated the offline recording session for these subjects to be able to retrain a classifier that achieved a performance significantly above random. S10 reported not being focused on Day-1. S7 and S9 were not available for the 3rd day of recording, therefore, they did not performed any online session. Table 2 summarizes the details of the recording sessions for each subject.

**Table 2.** Details of the recording sessions of Experiment-1.

| Subjects | # of offline sessions | # of online sessions |
| --- | --- | --- |
| S1, S2, S3, S4, & S5 | 1 | 2 |
| S6, S8 | 1 | 1 |
| S10 | 2 | 1 |
| S7, S9 | 2 | 0 |

During the online sessions, in addition to RT values (behavioral feedback), the output of real-time EEG classification was also provided to the subject (BCI feedback), which informed the subject whether or not the BCI could detect his/her intention to brake (i.e., presence of anticipatory SCPs before the onset of the ‘Red’). This was shown 2s after braking as a text appearing at the center of the screen which was ‘Yes’, if the BCI detected the movement intention, and ‘No’ otherwise. We hypothesized that providing the RT values as behavioral feedback to the subjects could help them better synchronize their actions (pressing the brake pedal) with the onset of the imperative cue (‘Red’).

To investigate the effect of providing behavioral and BCI feedback on the subject’s anticipatory behavior and its corresponding neural correlates, the evolution of EMG onset and the peak negativity at the Cz electrode were evaluated across different days. To estimate the peak negativity, we computed the minimum value within a 50 ms window from [-250, −200] ms with respect to the onset of the ‘Red’ light. These negativity values are obtained over each session for *Drive* and *Brake* trials separately. The temporal evolution of this negativity over a session was approximated with a linear polynomial. The slope of these lines is compared across days. The EMG onset distribution is defined as the time when the EMG activity exceeds a threshold equal to μ+ 6σ^13,17^, where *μ*and σ are the mean and standard deviation of the EMG in the window [-2.5, −2]s, during the timing of the *No-go* epoch (i.e., during the ‘Yellow’ light).

#### 2.1.2 Pre-processing

##### Offline sessions

The EEG data processing is similar to that of our previous study^13^. We first down-sampled the raw signals to 256 Hz, and discarded trials whose maximum potential exceeded 100 *μV*, then spatially filtered using a Common Average Reference (CAR)^19^. Next, the signal was smoothened using a Weighted Average filter (WAVG) applied to the CAR referenced data. WAVG can be seen as the opposite of the Laplacian filter^19^, where a channel’s average neighboring activity is added to it, rather than subtracted. Given the value of the *i^th^* electrode, *e_i_*(*t*) after CAR 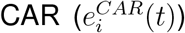, WAVG returns 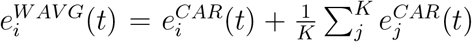, where *K* represents the number of nearest neigh bor electrodes considered (K=4 in this case, which all are equally weighted). It has been previously shown that such filtering improves the classification performance of CNV potentials^9^. The reason is that SCPs are widely spread over the central scalp areas. This means that similar patterns of SCPs are observed in the central electrodes, which can be taken into account in order to improve the Signal to Noise Ratio (SNR) using a weighted technique. Afterwards, the signals were spectrally filtered by means of a narrow bandpass Infinite Impulse Response (IIR) filter (4th order Butterworth in [0.1–1] Hz).

The EMG signals (down-sampled to 256 Hz) were filtered with a *bandpass* Butterworth filter in [20–50] Hz, then rectified and smoothened with a moving average filter (time window = 25 samples)^17^. Finally, the EEG and EMG signals were segmented into *Go*, and *No-go* epochs (2 s, starting at the onset of the warning stimulus). For each epoch the data were baseline-corrected by removing the value of the sample at the cue onset.

##### Online sessions

The processing steps for the real-time analysis of the online experiment were similar to the offline analysis, with the exception of the spectral filtering. The prerequisite for online experiments is to use causal filters (that can lead to significant changes on the signal morphology^18^), which may affect the classification performance. Thus, we compared the post-hoc analysis (offline classification performance) on the data of Day-1, for two different cases: with *bandpass* filters of [0.1–1] Hz applied causally and without the spectral filtering. An initial evaluation using the data recorded during the offline sessions yielded better training performance for the case with no spectral filter, and thus this setting was used in the online experiment.

#### 2.1.3 Feature extraction and classification

For the single-trial analysis, based on previous works^9,13^, features from the Cz electrode were extracted (see Figure 4.a). For each epoch, the Cz potentials at 4 equally spaced time points (−1.6 s, −1.2 s, −0.8 s, and −0.4 s) were used as a feature vector. This choice of the time points allowed online detection of the intention to act before subjects pressed the pedals. For classification, we used the Quadratic Discriminant Analysis (QDA)^8^ to discriminate between the *Go* and *No-go* epochs. Offline performances were assessed using a 4-fold cross-validation method that kept the chronological order of the data; i.e., each fold corresponds to a separate run. The performance of the single trial classification was evaluated using AUC in the ROC space^8^. ROC curves show the trade-off between the False Positive Rates (*FPR*) and True Positive Rates (*TPR*) of the classifier for different decision thresholds. In our case, *TPR* is the portion of *Go* epochs that are classified as *Go* and *FPR* is portion of *No-go* epochs erroneously detected as *Go* epochs. Online performance is reported in terms of the classifier accuracy.

### 2.2 Experiment-2: Real car

#### 2.2.1 Setup and protocol

Eight healthy subjects participated in the recordings comprising two sessions on took place within a period of 10 days (one subject could not join for Day-2). None of the subjects in this experiment took part in Experiment-1 in the car simulator. Experiment-2 was performed with a real automobile with automatic gear (*Infinity FX30*, see Figure 3.a). For safety reasons, the experiments were performed in a closed road with no other vehicle or pedestrians present (Figure 3.b). All experiments were done in daylight, spanning 10 consecutive days and under similar weather conditions (cloudy/sunny). In this road, we placed 6 traffic lights at specific locations (see Figure 3.c). During the experiments all driving assistance systems such as intelligent cruise control were disabled. Moreover, the drivers were instructed to use the automatic gearshift and to keep their hand on the steering wheel to limit arm movements.

**Figure 3:**
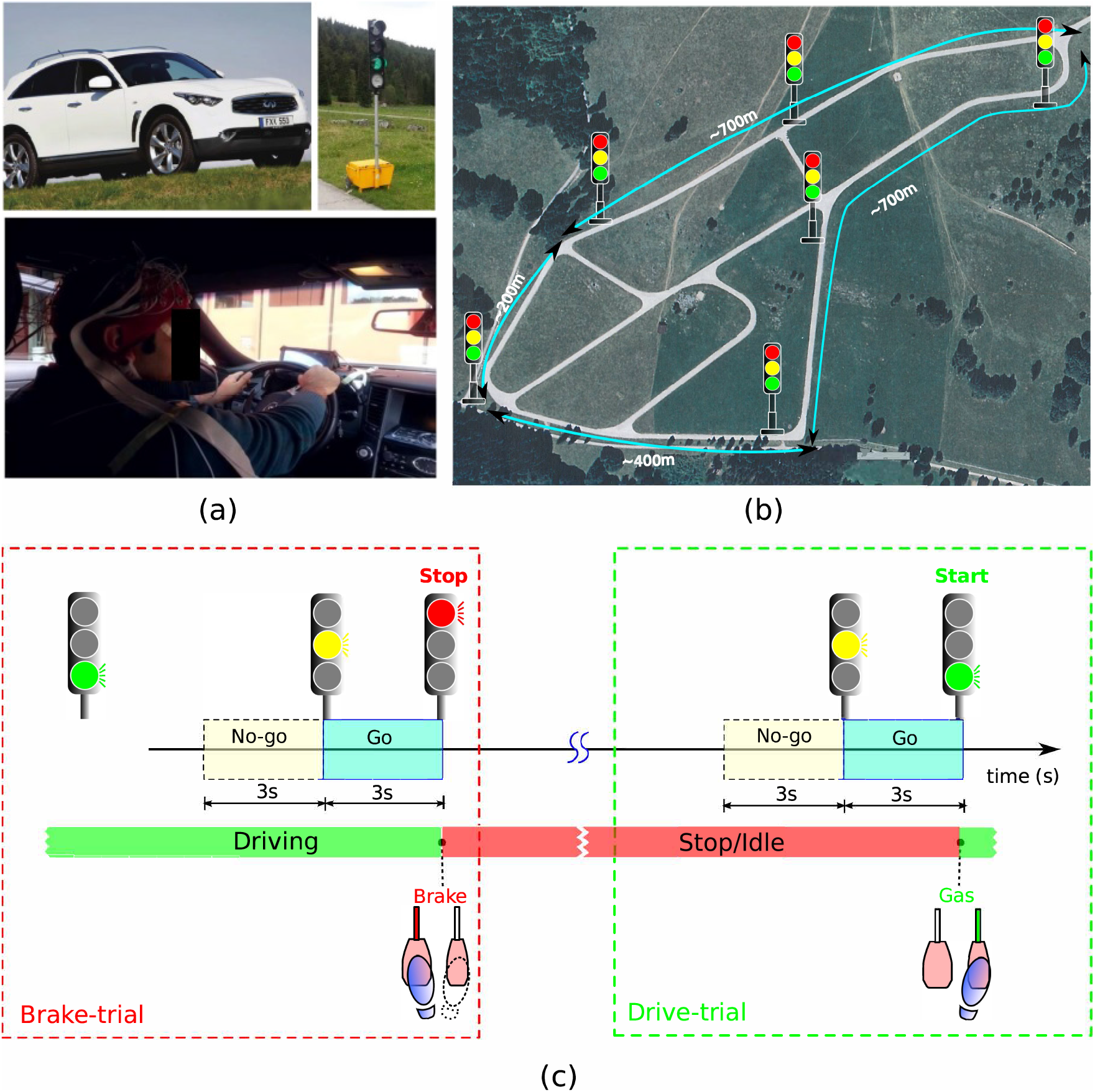
Experiment-2 setup and timeline of the trials, (a) The vehicle Infiniti FX30, and the subject in the car, the real traffic lights used in the experiment. (b) Aerial view of the road and location of the traffic lights. (c) Time-line of the protocol. The traffic lights turning from ‘Green’ to ‘Yellow’ to ‘Red’ followed by braking after the ‘Red’ light are called *Brake* trials. ‘Red’ to ‘Yellow’ to ‘Green’ correspond to *Drive* trials. Each trial contains one *Go* and one *No-go* epoch.

These traffic lights were programmed with fixed timings (‘Green’: 10 s ‘Yellow’: 3 s,’Red’: 27 s). Subjects were asked to drive normally at about 60 Km/h so that the car would arrive at each traffic light location at specific times that yielded *Go* and *No-go* epochs of 3 s. As in the previous experiment, we defined two types of trials: *Drive* and *Brake* (see Figure 3.c). The experimental design allowed us to have 4 *Brake* and 6 *Drive* trials per lap. Each recording session consisted of five laps driving along the circuit, which lasted around 15 to 20 minutes, depending on the driving style of each subject.

An average (across all the subjects and all the sessions) of 64±15 trials for *Drive* and 43±11 for *Brake* trials were obtained. The data were labeled manually by the experimenter, sitting in the same car. Limited availability to the closed road did not allow to perform enough recording sessions to perform an online test for this experiment.

#### 2.2.2 Pre-processing, Feature extraction, and classification

Similar signal processing steps used for experiment-1 have been explored for Experiment-2. The few differences are described below: For the analysis, we selected the central electrodes (C1, C2, Cz, FC1, FCz, FC2, CP1, CPz, and CP2), as the outer electrodes were more prone to contamination by driving related artifacts (eye and neck movements and EEG cap touching the headrest of the driver seat). Given the reduced montage, for the spatial filtering we avoided using CAR as it would reduce anticipatory SCP negativity in electrodes of interest (central electrodes). Thus, we directly applied the WAVG smoothing filter introduced in the section 2.1.2. Due to the timing of the traffic lights, the Inter-Stimulus-Interval in this experiment is 3s (see Figure 3.c). However, we used the same features as for Experiment-1; i.e., 4 equally spaced time-points extracted from the last 2s before each cue. This choice was based on the analysis of EEG grand averages from the data recorded on Day-1, showing that the negative slope starts around 1 s after the ‘Yellow’ light. The classifiers and the methods for evaluating the classification performance were the same as for offline sessions of Experiment-1.

## 3 Results

### 3.1 Experiment-1: EEG correlates of anticipation in the car simulator

#### 3.1.1 Grand averages

Figure 4 illustrates the EEG grand averages for the *Drive* and *Brake* trials, computed across all subjects on the recordings of Day-1. As shown on the scalp maps at different time points (top panel of Figure 4), the negative potential is evident in the central area and is maximal at centro-medial electrodes. This observation is consistent with results reported in existing literature on anticipation-related SCPs^27^ as well as in our previous work^13^.

**Figure 4:**
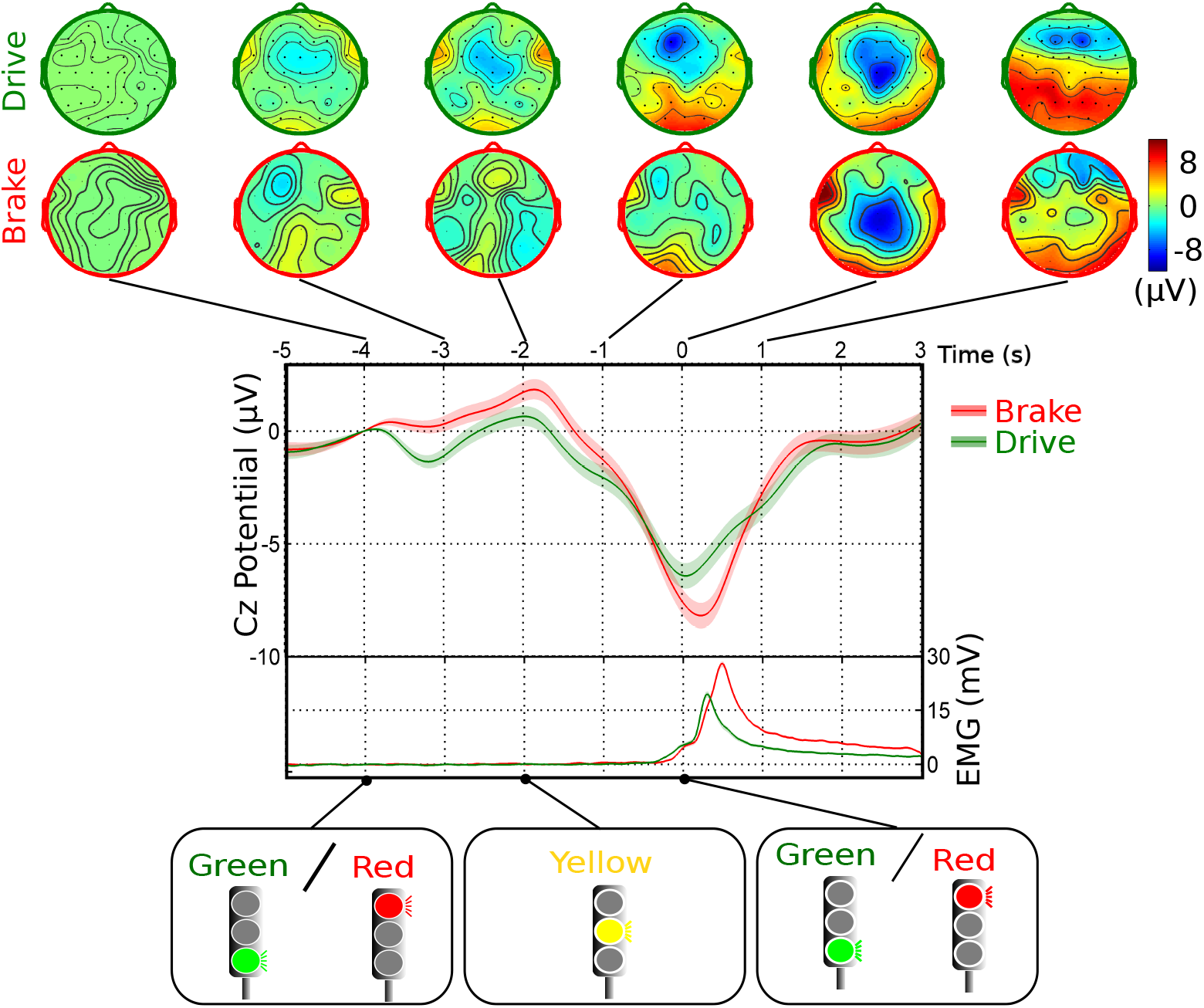
Grand averages of Cz potentials for the Experiment-1 in the car simulator on Day-1. (top) Topographic representation of average EEG scalp distribution at different time points. (bottom) Grand averages of Cz potentials and EMG envelopes shown for *Drive* (in green) and for *Brake* trials (in red). The mean is shown with solid line and the standard error shown in shadow (for Cz potentials it is < 1.5μV and for EMG it is <1.5 mV). At time t=0s is the onset of the appearance of ‘Green’/’Red’ light (imperative stimulus). EEG and EMG data were baseline corrected to the value of the sample at the onset of ‘Red’/’Green’ light at t=-4s.

A negative amplitude deflection at the Cz electrode starting at the onset of the ‘Yellow’ light can be observed for both *Drive* and *Brake* trials, reaching its maximum negativity slightly after t=0s. The time point t=0s, corresponds to the onset of ‘Green’/’Red’. As shown in Figure 4, a higher average peak negativity appears for *Brake* than for *Drive*trials (Wilcoxon test, p-value <0.01). The bottom panel of Figure 4 show the grand averages of the EMG envelopes, demonstrating that there is no muscular activity during the preparation phase.

#### 3.1.2 Single-trial classification

Figure 5 shows the ROC curves of individual offline classification performance (4-fold cross-validation) in the car simulator for *Drive* and *Brake* trials, separately. AUC above 0.80 were obtained for five subjects, in at least one of the sessions for *Brake* (S2, S4, S5, S6, and S8) and for *Drive* trials (S3, S4, S5, S8, and S9). The mean AUC across 10 subjects for Day-1 was 0.73±0.15 for *Drive* trials and 0.73±0.14 for *Brake* trials. On Day-2, the mean AUC was 0.67±0.14 for *Drive* trials and 0.73±0.14 for *Brake* trials. For Day-3, the mean AUC across the 6 subjects that participated in the third day were 0.73±0.09 and 0.80±0.08 for *Drive* and *Brake* trials, respectively. Thus, we observe that the average classification performance for *Brake* trials is superior to that of *Drive* trials throughout the 3 days of experiment in the car simulator and the 2 days with the real car, but their differences are not statistically significant (Wilcoxon test).

**Figure 5:**
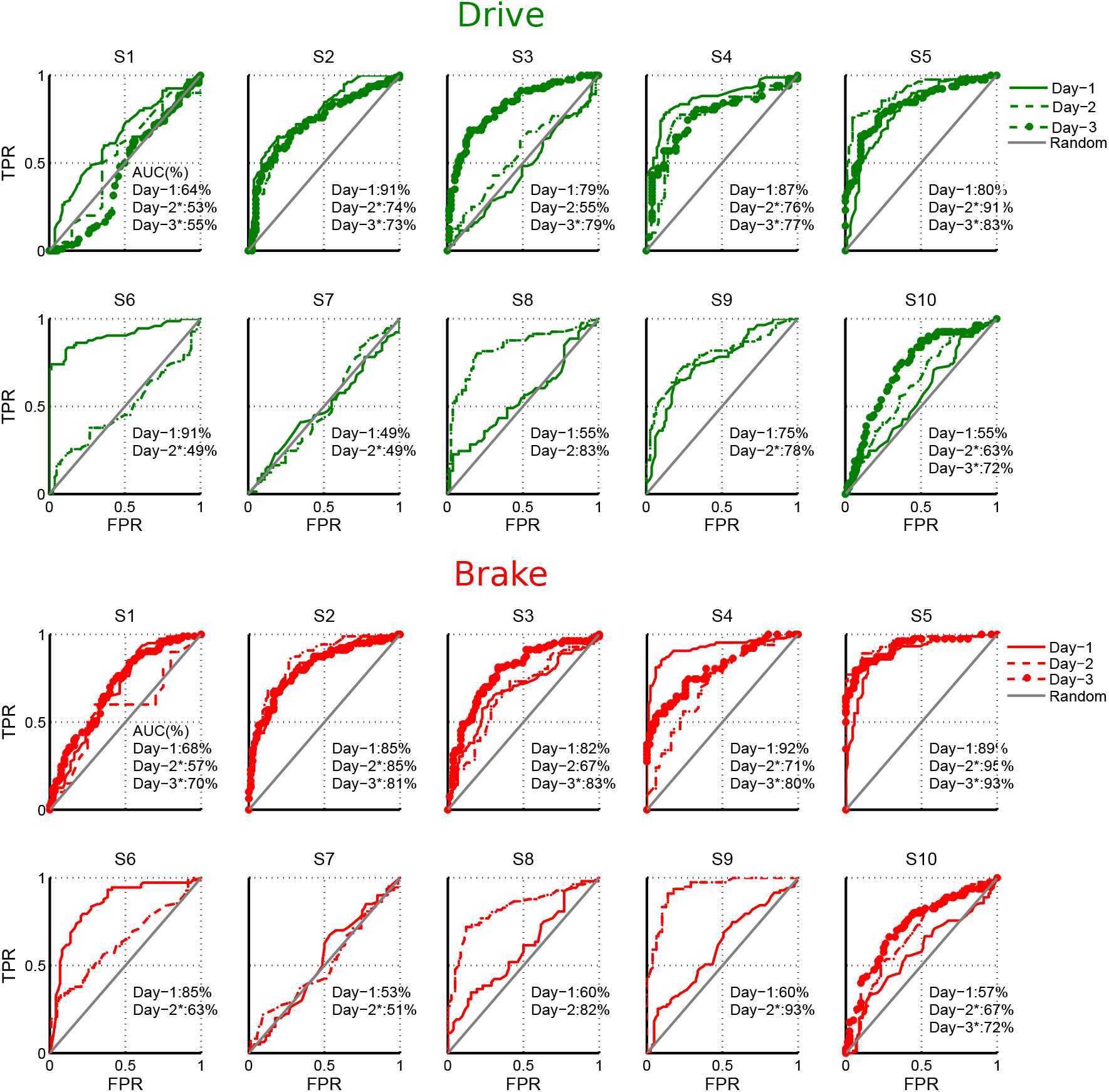
Experiment-1. Individual classification performance (AUC) in the car simulator for *Drive* (in green) and *Brake* trials (in red). ROC curves and mean AUC values for all subjects on different days of recording. The gray diagonal line represents random performance.

The performance of the online experiment for *Brake* trials was evaluated as the accuracy of the real-time classification results of the *Go* and *No-go* epochs (see Table 3). Noteworthy, the accuracy exceeded 0.60 for 5 subjects, which corresponds to the 95% confidence interval of chance level for a two-class problem when about 80 trials per class are available^21^. Moreover, two out of the 3 subjects who did not reach this threshold in Day-2, did in Day-3.

**Table 3.** Online classification accuracies in the Experiment-1.

| | S1 | S2 | S3 | S4 | S5 | S6 | S8 | S10 | Mean±SD |
| --- | --- | --- | --- | --- | --- | --- | --- | --- | --- |
| Day-2 | 0.58 | 0.51 | 0.52 | **0.76** | **0.75** | 0.50 | **0.83** | — | 0.63±0.13 |
| Day-3 | 0.55 | **0.66** | **0.72** | **0.66** | **0.69** | — | — | **0.61** | 0.65±0.05 |

#### 3.1.3 Effect of feedback

We report changes in the response time and the SCPs measured from 5 subjects (S1 to S5) for whom we had measurements over all three days; Day-1 (offline): RT feedback for *Brake* trials; Day-2 (online) and Day-3 (online): RT+BCI feedback for *Brake* trials (Table 1 and 2). To investigate the effect of feedback we first computed the EMG onset timing, which representing the subject’s response time. We then computed the median of the EMG onset time of Day-1 as a reference. Following this, we computed the percentage of trials that have EMG onset time above this value for Day-2 and Day-3 (shown in Figure 6.A).

**Figure 6:**
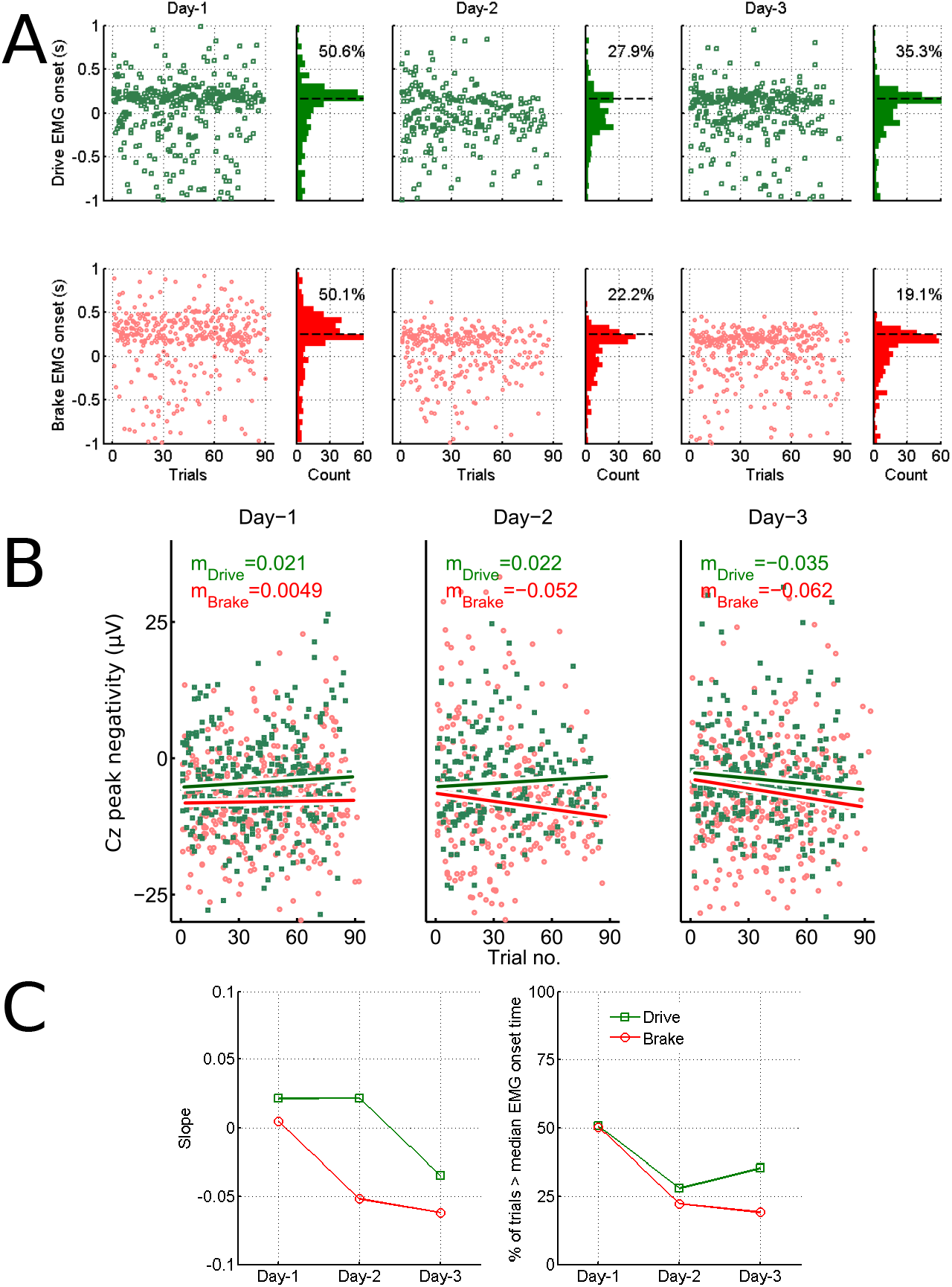
A)The evolution of EMG onset time for the 5 subjects (S1, S2, S3, S4, S5) for *Drive* (in green) and *Brake* for the 3 recording days. The dotted line corresponds to median value of Day-1. The number in the distribution plot describes the percentage of trials with EMG onset time above the median value of Day-1. B) The peak negativity value of Cz potentials for *Drive* (in green) and *Brake* (in red). The slope of a line fitted to the peak negativity across trials is also displayed. C) Summary for the 3 days: (Left) The slope of the fitting line. (Right) The percentage of trials above the median EMG onset.

Generally, the median of the EMG onset timing is higher for *Brake* trials (246 ms, 152 ms and 160 ms) than for *Drive* trials (156 ms, 98 ms, 121 ms) for Day-1, Day-2, and Day-3, respectively. This likely due to the need for a more complex movement for braking. Here the subjects need to switch their foot from the gas pedal to the brake, whereas during the *Drive* the subject’s foot was already on the gas pedal.

We compare the evolution of the EMG onset timing across different days for *Drive* and *Brake* trials, as the former receive no feedback in all the 3 days and the latter receive the RT feedback on Day-1 (plus BCI feedback on Day-2 and Day-3). Indeed, we observe that subjects become more predictive of the imperative stimulus (i.e., appearance of ‘Green’ or ‘Red’) rather than reactive from Day-1 to Day-2 and Day-3. The results show that, for *Drive*, the percentage of trials above the median of the EMG onset distribution of Day-1 (=156 ms), was 50.6% and reduced to 27.9% and 35.3% during Day-2 and Day-3, respectively. Similarly, for *Brake*, the percentage of trials above the median EMG distribution of Day-1 (=246 ms) was 50.1% and reduced to 22.2% and 19.1% for Day-2 and Day-3, respectively.

Reduction in the EMG onset time for *Brake* trials is likely to be a signature of an improvement in anticipatory behavior, which could be due to the feedback (RT+BCI) provided on Day-2 and Day-3. We observe a consistent reduction of this value across days for Brake trials as compared to *Drive* trials. However, from the current data it is not possible to dissociate the contribution of each of these feedback components to the learning effect (note that the sessions of RT alone and RT+BCI feedback are not randomized).

Since BCI feedback is the result of the real-time classification of the centromedial EEG negativity, we investigated the evolution of this negativity throughout each experimental session across the different recording days. The change in the negativity of the SCP potentials within a session was expressed as the slope (= change in peak negativity across different trials) of a line fitting the peak negativity values within a session, as shown in Figure 6.B. The slopes of the *Drive* trials were 0.021, 0.022 and −0.035 *μ*Vs per trial over Day-1, Day-2 and Day-3, respectively. For the *Brake* trials, the slopes were 0.0049, −0.052 and −0.062 *μ*Vs per trial over Day-1, Day-2 and Day-3, respectively. We observe that the negativity mainly increased in *Brake* (Day-2 and Day-3) trials as compared to *Drive* trials (only Day-3). The slope is close to zero for *Brake* of Day-1, when only RT feedback was provided, indicating no change in negativity across trials. A nonzero negative slope is observed for trials of Day-2 and Day-3, when BCI feedback was also delivered, that could suggest a decrease of the Cz peak value. Hence, further investigation is needed to assess this tendency.

### 3.2 Experiment-2: EEG correlates of anticipation in real car

#### 3.2.1 Grand averages

Figure 7 illustrates the grand averages of *Drive* and *Brake* trials in the real car. The neural signatures are very similar to those observed in the car simulator experiment. The negativity for *Brake* trials starts around 2s before the appearance of the ‘Red’; (around 1 s after the onset of ‘Yellow’). This negativity starts around the same timing for the *Drive*, initially with a smaller slope then with a bigger slope around 1 s before the onset of ‘Green’ light. The topographic plots of the real car experiment reveal an average negativity localized in the central area, similar to what was observed previously. Regarding EMG, their envelop shows that the onset of increasing EMG activity occurs before 0s. The time point t=0s, corresponds to the onset of ‘Green’/’Red’, as manually marked by the experimenter.

**Figure 7:**
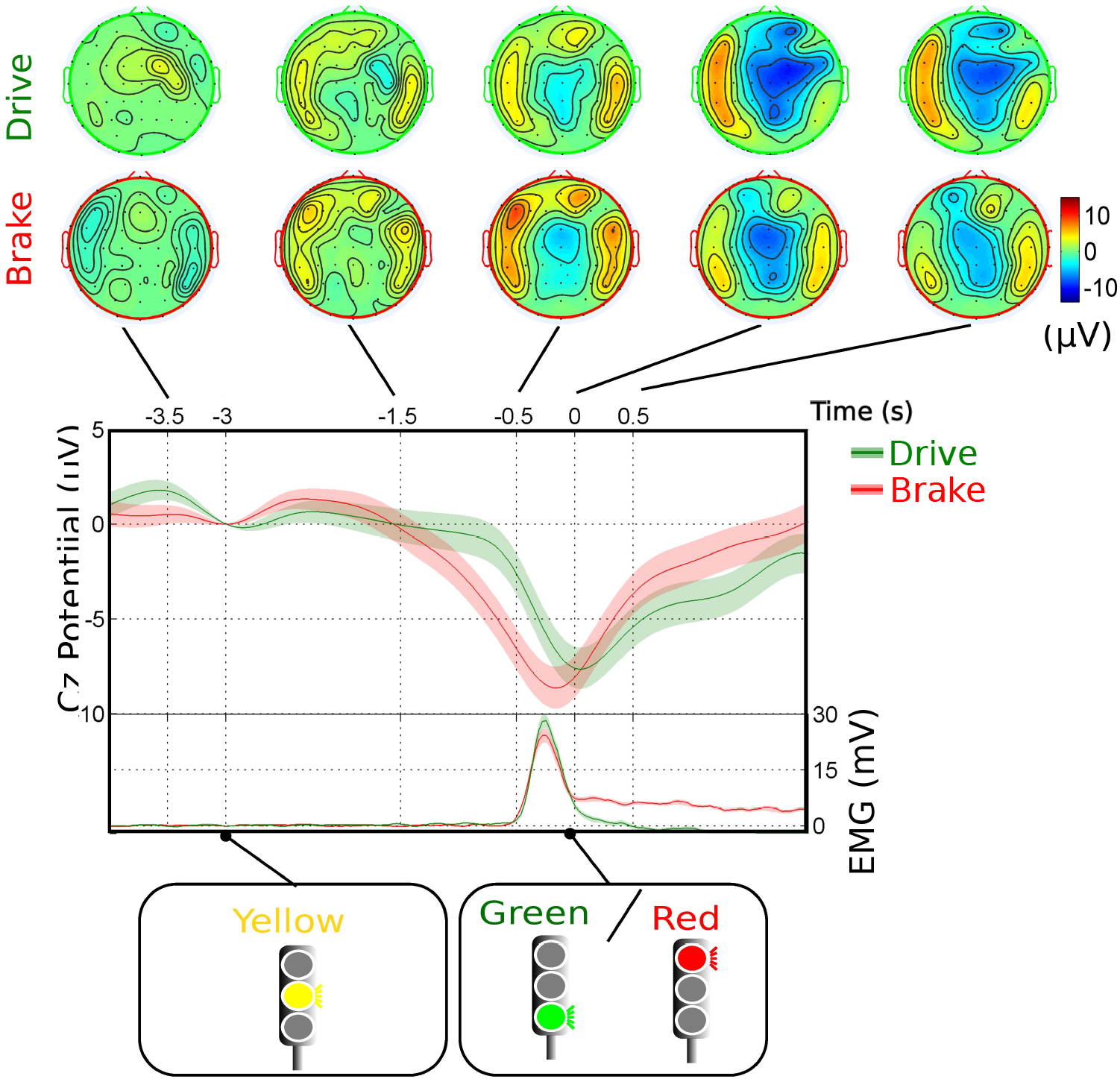
Grand averages of Cz potentials for the real car experiment on Day-1. (top) Topographic representation of average EEG scalp distribution at different time points. (bottom) Grand averages of Cz potentials and EMG envelopes shown for *Drive* (in green) and for *Brake* (in red). The mean is shown with solid line and the standard error shown in shadow. t=0 s is the onset of ‘Green’/’Red’ and, t=-3s is the onset of ‘Yellow’. Data were baseline corrected to the value of the sample at the onset of ‘Yellow’.

#### 3.2.2 Single-trial classification

Figure 8 illustrates the individual single-trial classification of Cz potential for *Drive* and *Brake* trials. The mean AUC of the *Drive* trials were 0.63±0.08 and 0.62±0.07 for Day-1 and Day-2, respectively. The mean AUC of the *Brake* trials were 0.66±0.13 and 0.63±0.12 for Day-1 and Day-2, respectively. Notably, 4 subjects for *Drive* trials and 4 subjects for *Brake* trials reached an AUC of 0.70. The reason for the odd behavior of the ROC curves for S17 on Day-2 could be the very limited number of trials for these subjects in order to estimating the parameters of the classifier.

**Figure 8:**
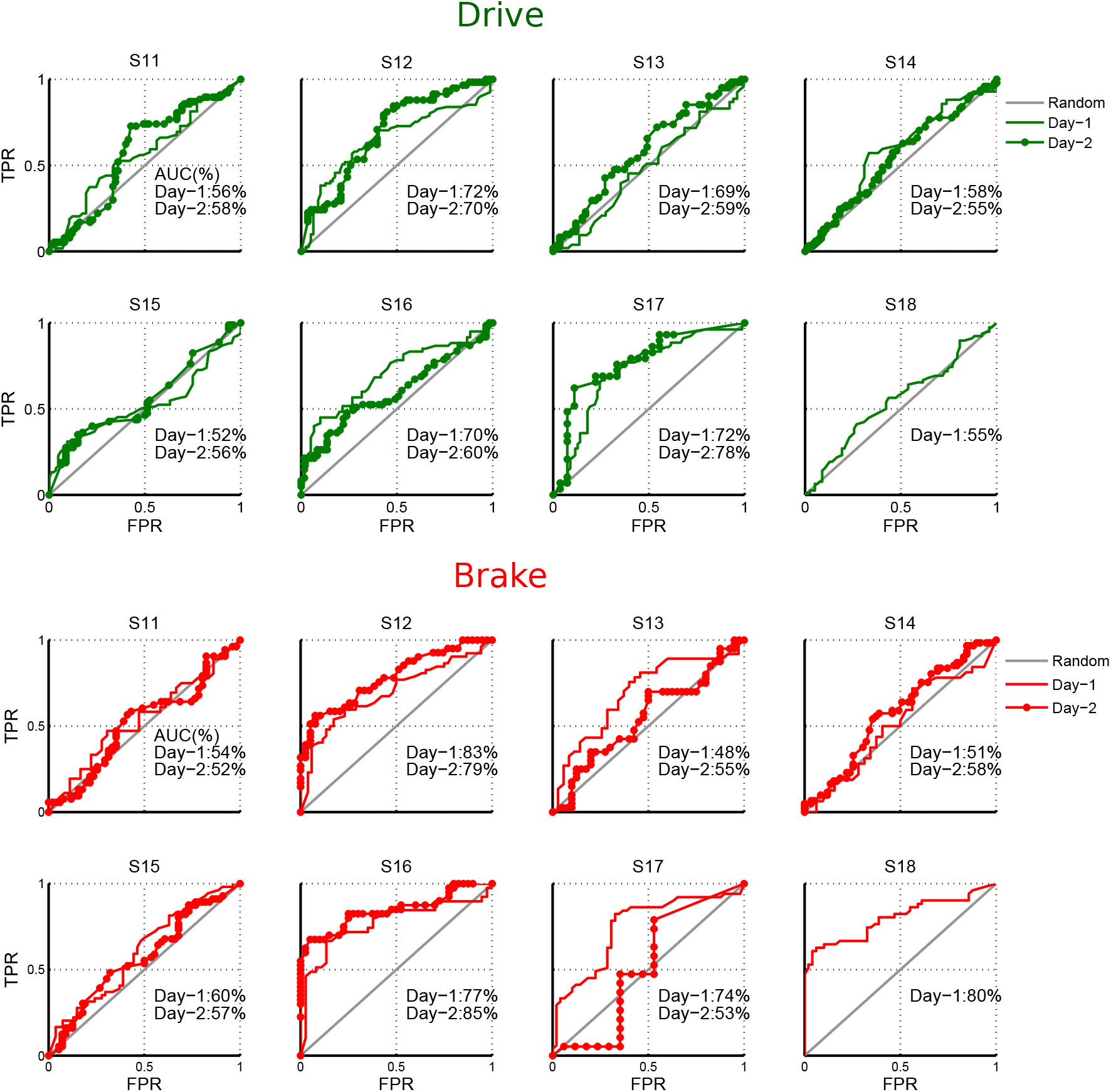
ROC curves showing the classification performance of different subjects for *Drive* (in green) and *Brake* (in red) of the data recorded from Experiment-2 (with real car). The AUC values on different recording days are displayed.

## 4 Discussion

This paper extends our previous work^13^ on investigating the neural correlates of anticipatory behavior to more realistic driving scenarios analyzing this type of behavior in response to traffic light changes. These scenarios include simulated and real-world driving. Firstly, we proposed a realistic simulated scenario, recording 10 subjects, to investigate the anticipatory brain potentials in a virtual city with several junctions and traffic lights. We demonstrated the presence of anticipatory SCPs in response to traffic lights with similar patterns as those observed in our previous work where a count-down stimuli was used^13^ as well as the classical CNV signals reported in the literature^27,9^.

The single-trial classification analysis revealed satisfactory performances (in BCI decoding). Moreover, we tested the possibility of online decoding of the intention to brake in the car simulator, which yielded accuracy levels significantly above chance (average: 0.64±0.1). All subjects who did two online sessions had better performance in the last day. It has been suggested that closing the loop enhances BCI performances by exploiting the learnability potentials of the human brain. ^28,23^ Our preliminary findings suggest that providing the result of the real-time classification of EEG together with the behavioral feedback can lead to subjects learning of anticipatory behavior.

We further extended the experimental protocols to a real car while driving on a closed road, recording 8 subjects. Remarkably, anticipatory SCPs were also observed for the first time in this scenario and resembled those of the car simulator study. However a significant decrease is observed in the classification performance of the real car data in comparison to the car simulator study (Wilcoxon test, p-value=0.01). This is probably due to the noisier signals (due to artifacts and the complexity of the task) during the real-world driving.

Arguably, the anticipatory behavior may be affected by visual distractions and multitasking nature of motor control in driving. Thus, these artifacts and noise are likely to degrade the quality of the measured brain signals in real-world driving settings. It is worth noting that the experimenter manually tagged events in the real car experiment (the experimenter was seatted in the backseat of the car and manually entered the events using a button sampled together with EEG and EMG), which has inevitably introduced considerable jitters. Despite this jitter, it is indeed encouraging that SCPs seem to be sufficiently robust to appear in grand averages and be detectable at the single-trial level. Future experiments can be implemented with an automatic labeling system, where in-car embedded sensors would be used for detecting the traffic light changes^6,7,22^, which could be transferred to the proposed BCI system. This approach may minimize the jitters and hence increase SNR.

The main goal of this work was to assess the feasibility of reliable detection of anticipatory SCPs in real-time during real world driving, in spite of enriched visual distractions, and numerous sources of artifacts is the main contribution of this work. For that, one needs to identify the common sources of the noise and artifacts during driving and accordingly apply pre-processing methods for their removal. Additionally driving related movement artifacts could be reduced through Independent Component Analysis (ICA) or linear regression models^5,24^.

We have performed the experiments in both car simulator and real car on the normal allocated road. In the future, it is also necessary to perform similar assessments in real traffic conditions, ranging from the inclusion of other distracting elements such as multiple automobiles and pedestrians to the driver could drive the car at a variable speed that suits the traffic conditions.

Finally, another current limitation of the proposed BCI for smart cars regards the need for large number of electrodes (mainly for the spatial filtering) what leads to an expensive, obtrusive, and not easy to setup system. Developing an in-car BCI system requires a reduced number of electrodes. Alternatively, methods for choosing only the relevant and discriminant channels from a large number of channels (in our case 64) have been already proposed in other EEG studies^16,20,25^ and recently also for the recognition of anticipation related potentials^9^. Furthermore, ‘dry’ electrodes have been developed to increase ease of use^10^. Preliminary tests with a dry electrodes systems (not reported in this paper) suggest that montages as small as 16 channels yield similar performances as those reported here using the 64 channels.

In conclusion, the main contribution of this paper is the exploration of anticipatory behavior in simulated and real car driving, where in both cases, we report the presence of anticipatory SCPs, similar to the well known CNV potentials. Furthermore, we report an online detection system that provided BCI feedback. These findings can be beneficial in detecting the planned action before its execution during driving.

## Acknowledgment

This study was supported by Nissan Motor Co. Ltd., and the Swiss-funded National Centre of Competence in Research (NCCR) Robotics. The authors would like to thank all participants, and acknowledge the valuable assistance and support from Michael Themans (EPFL transportation center, Lausanne Switzerland) and Jonathan Martin (Cantonal road service, Vaud, Switzerland). Real car experiments were possible thanks to the support of the Cantonal road service of Vaud, Switzerland.

## References

1. https://www.blender.org/download/.

2. Birbaumer, N., T. Elbert, A. G. Canavan, and B. Rockstroh. Slow potentials of the cerebral cortex and behavior. Physiological reviews 70:1–41, 1990.

3. Blankertz, B., M. Tangermann, C. Vidaurre, S. Fazli, C. Sannelli, S. Haufe, C. Maeder, L. Ramsey, I. Sturm, G. Curio et al. The Berlin brain–computer interface: non-medical uses of BCI technology. Frontiers in neuroscience 4, 2010.

4. Chai, R., S. H. Ling, P. P. San, G. R. Naik, T. N. Nguyen, Y. Tran, A. Craig, and H. T. Nguyen. Improving eeg-based driver fatigue classification using sparse-deep belief networks. Frontiers in neuroscience 11, 2017.

5. Daly, I., R. Scherer, M. Billinger, and G. Müller-Putz. Force: Fully online and automated artifact removal for brain-computer interfacing. IEEE transactions on neural systems and rehabilitation engineering 23:725–736, 2015.

6. De Charette, R. and F. Nashashibi. Real time visual traffic lights recognition based on spot light detection and adaptive traffic lights templates. In: IEEE Intelligent Vehicles Symposium, pp. 358–363. 2009.

7. Diaz, M., P. Cerri, G. Pirlo, M. A. Ferrer, and D. Impedovo. A Survey on Traffic Light Detection, pp. 201–208, Cham: Springer International Publishing 2015.

8. Duda, R. O., P. E. Hart, and D. G. Stork. Pattern Classification, New York: Wiley 2001, edition.

9. Garipelli, G., R. Chavarriaga, and J. d. R. Millán. Single trial analysis of slow cortical potentials: A study on anticipation related potentials. Journal of Neural Engineering 10:036014, 2013.

10. Guger, C., G. Krausz, and G. Edlinger. Brain-computer interface control with dry EEG electrodes. 2011.

11. Haufe, S., J. Kim, I. Kim, A. Sonnleitner, M. Schrauf, G. Curio, and B. Blankertz. Electrophysiology-based detection of emergency braking intention in real-world driving. Journal of Neural Engineering 11:056011, 2014.

12. Hayakawa, Y., K. Sato, Y. Tabata, and K. Egawa. Design of a lane departure prevention system with enhanced drivability. SAE International Journal of Passenger Cars-Mechanical Systems 2:398–403, 2009.

13. Khaliliardali, Z., R. Chavarriaga, L. A. Gheorghe, and J. d. R. Millán. Action prediction based on anticipatory brain potentials during simulated driving. Journal of Neural Engineering 12:066006, 2015.

14. Kim, I. H., J. W. Kim, S. Haufe, and S. W. Lee. Detection of braking intention in diverse situations during simulated driving based on EEG feature combination. Journal of Neural Engineering 12:016001, 2015.

15. Knight, J. Vdrift open source drift racing simulator, 2008. http://www.linuxjournal.com/article/10214, accessed 2008-10-01.

16. Lan, T., D. Erdogmus, A. Adami, M. Pavel, and S. Mathan. Salient EEG channel selection in brain computer interfaces by mutual information maximization. In: Engineering in Medicine and Biology Society, 2005. IEEE-EMBS 2005. 27th Annual International Conference of the, pp. 7064–7067.

17. Lew, E., R. Chavarriaga, S. Silvoni, and J. d. R. Millán. Detection of self-paced reaching movement intention from EEG signals. Frontiers in Neuroengineering 5:13, 2012.

18. Luck, S. J. An introduction to the event-related potential technique, MIT press 2014.

19. McFarland, D. J., L. M. McCane, S. V. David, and J. R. Wolpaw. Spatial filter selection for EEG-based communication. Electroencephalography and Clinical Neurophysiology 103:386–394, 1997.

20. Millán, J. d. R., M. Franzé, J. Mouriño, F. Cincotti, and F. Babiloni. Relevant EEG features for the classification of spontaneous motor-related tasks. Biological cybernetics 86:89–95, 2002.

21. Müeller-Putz, G., R. Scherer, C. Brunner, R. Leeb, and G. Pfurtscheller. Better than random: A closer look on BCI results. International Journal of Bioelectromagnetism 10:52–55, 2008.

22. Omachi, M. and S. Omachi. Traffic light detection with color and edge information. In: 2nd IEEE International Conference on Computer Science and Information Technology, pp. 284–287, IEEE 2009.

23. Perdikis, S., R. Leeb, and J. d. R. Millán. Subject-oriented training for motor imagery brain-computer interfaces. In: Engineering in Medicine and Biology Society (EMBC), 2014 36th Annual International Conference of the IEEE, pp. 1259–1262, IEEE 2014.

24. Schlögl, A., C. Keinrath, D. Zimmermann, R. Scherer, R. Leeb, and G. Pfurtscheller. A fully automated correction method of EOG artifacts in EEG recordings. Clinical Neurophysiology 118:98–104, 2007.

25. Schröder, M., T. N. Lal, T. Hinterberger, M. Bogdan, N. J. Hill, N. Birbaumer, W. Rosenstiel, and B. Schölkopf. Robust EEG channel selection across subjects for brain-computer interfaces. EURASIP Journal on Applied Signal Processing 2005:3103–3112, 2005.

26. Takae, Y., Y. Seto, T. Yamamura, T. Sugano, M. Kobayashi, and K. Sato. Development and evaluation of a distance control assist system with an active accelerator pedal. Technical report, SAE Technical Paper, 2009.

27. Walter, W. G., R. Cooper, V. J. Aldridge, and W. C. Mccallum. Contingent negative variation : An electric sign of sensorimotor association and expectancy in the human brain. Nature pp. 380–384, 1964.

28. Wander, J. D., T. Blakely, K. J. Miller, K. E. Weaver, L. A. Johnson, J. D. Olson, E. E. Fetz, R. P. Rao, and J. G. Ojemann. Distributed cortical adaptation during learning of a brain–computer interface task. Proceedings of the National Academy of Sciences 110:10818–10823, 2013.

29. Zhang, H., R. Chavarriaga, Z. Khaliliardali, L. A. Gheorghe, and J. d. R. Millán. EEG-based decoding of error-related brain activity in a real-world driving task. Journal of Neural Engineering 12:066028, 2015.

